# Müller glial Kir4.1 channel Dysfunction in *APOE4*-KI model of Alzheimer’s disease

**DOI:** 10.1101/2025.02.26.640427

**Authors:** Surabhi D. Abhyankar, Yucheng Xiao, Neha Mahajan, Qianyi Luo, Theodore R. Cummins, Adrian L. Oblak, Bruce T. Lamb, Timothy W. Corson, Ashay D. Bhatwadekar

**Author notes:** Correspondence: Ashay Bhatwadekar, PhD, RPh, FARVO, Department of Ophthalmology, 1160 W Michigan Street, GK-305P, Indianapolis, IN-46202. **Email addresses:** Surabhi D. Abhyankar, Yucheng Xiao, Neha Mahajan, Qianyi Luo, Theodore R. Cummins, Adrian L. Oblak, Bruce T. Lamb, Timothy W. Corson.

## Abstract

Alzheimer’s disease (AD), particularly late-onset AD (LOAD), affects millions worldwide, with the apolipoprotein ε4 (*APOE4*) allele being a significant genetic risk factor. Retinal abnormalities are a hallmark of LOAD, and our recent study demonstrated significant age-related retinal impairments in *APOE4*-knock-in (KI) mice, highlighting that retinal impairments occur before the onset of cognitive decline in these mice. Müller cells (MCs), key retinal glia, are vital for retinal health, and their dysfunction may contribute to retinal impairments seen in AD. MCs maintain potassium balance via specialized inwardly rectifying K^+^ channels 4.1 (Kir4.1). This study posits that Kir4.1 channels will be impaired in *APOE4*-*KI*, resulting in MC dysfunction. Additionally, we demonstrate that MC dysfunction in *APOE4*-*KI* stems from alterations in mitochondrial dynamics and oxidative stress. Kir4.1 expression and function were studied using immunofluorescence and through the whole-cell voltage clamp, respectively. In parallel, rat Müller cells (rMC-1) were used to create an *in vitro* model for further mechanistic studies. Mitoquinol (MitoQ) was used to evaluate its potential to mitigate *APOE4*-induced deficits. *APOE4* retinas and *APOE4*-transfected rMC-1 significantly reduced Kir4.1 expression, K+ buffering capacity, and increased mitochondrial damage. *APOE4*-transfected rMC-1 showed reduced mitochondrial membrane potential (ΔΨm) and increased mitochondrial reactive oxygen species (ROS). MitoQ treatment significantly reduced mitochondrial ROS and restored Kir4.1 expression in *APOE4*-expressing cells. Our results demonstrate that *APOE4* causes mitochondrial dysfunction and MC impairment, which may contribute to retinal pathology in AD. MitoQ restored mitochondrial health and Kir4.1 expression in *APOE4*-expressing rMC-1, suggesting targeting mitochondria may offer a promising therapeutic strategy for AD.

**Main Points:**

- *APOE4* impairs Müller cell health by reducing Kir4.1 expression and buffering.
- *APOE4* causes mitochondrial dysfunction with decreased ΔΨm and increased ROS.
- MitoQ restores Kir4.1 expression and reduces ROS in *APOE4*-transfected cells.

## 1 INTRODUCTION

Over 55 million people worldwide are living with dementia, with Alzheimer’s Disease (AD) being its most common form, responsible for roughly 60-80% of cases globally (World Health Organization Alzheimer’s Report, 2021). In the United States (US) alone, an estimated 6.7 million individuals aged 65 and older were diagnosed with AD in 2023. AD progressively impairs memory, learning, and executive functions (Bondi et al., 2017; Breijyeh & Karaman, 2020), making it increasingly difficult for individuals to make decisions, solve problems, communicate, or care for themselves (Silva et al., 2019). The most prevalent subtype of AD is late-onset AD (LOAD), which represents ∼95% of all AD cases worldwide (Boutajangout & Wısnıewskı, 2013) and affects nearly 30% of people over age 85 (Brodtmann, 2013; Darby, 2013). The *APOE4* allele is recognized as a significant genetic risk factor for LOAD (Uddin et al., 2018; Yamazaki et al., 2019), with 56% of AD patients in the US carrying one copy of the *APOE4* allele and 11% carrying two copies (“2023 Alzheimer’s disease facts and figures,” 2023; Rajan et al., 2021). The three *APOE* gene variants *APOE2* (cystine 112, cystine 158), *APOE3* (cystine 112, arginine 158) and *APOE4* (arginine 112, arginine 158) have differing effects on AD risk: while the *APOE4* allele raises risk, the *APOE2* allele is considered protective, and *APOE3* allele is neutral (Husain et al., 2021; Roses, 1996).

The retina shares many characteristics with the brain, including vascular connections, neural pathways, and immune regulations, and it often mirrors brain pathology (Golzan et al., 2017; Lim et al., 2016; Patton et al., 2005). Recent studies from our group have shown that 52-57-week-old *APOE4*-knock mice had retinal structural, functional, vascular, and vision deficits, increased neuroinflammation, and downregulation of synaptogenesis, suggesting middle-aged *APOE4* mice have retinal dysfunction (Abhyankar et al., 2025). Müller cells (MCs), the most abundant retinal glial cells, span the retina and provide structural support to neurons (Kobat & Turgut, 2020; Reichenbach & Bringmann, 2013), akin to astrocytes in the brain, helping to maintain the blood-retinal barrier by stimulating the production of tight junction proteins in endothelial cells (Bernardos et al., 2007). MC gliosis, a hallmark of AD-related pathology, involves generalized and potentially protective responses, such as elevated glial fibrillary acidic protein (GFAP) and diminished glutamine synthetase (GS) levels (Andreas Bringmann et al., 2006). Prominent MC activation has been observed in several AD mouse models, including App^NL-G-F^, 5xFAD, and 3xTG, emphasizing its significance in disease progression (Edwards et al., 2014; Vandenabeele et al., 2021; Zhang et al., 2021). Consistent with these findings, decreased GS levels have been reported in the brains (Kulijewicz-Nawrot et al., 2013; Le Prince et al., 1995; Olabarria et al., 2011; Robinson, 2001) and the retinas of individuals with AD (Tams et al., 2022; Xu et al., 2022).

MCs perform crucial roles in neurotransmitter uptake, glycogen storage, and maintaining water and K^+^ balance (A. Bringmann et al., 2006; Kobat & Turgut, 2020), largely through inwardly rectifying K^+^ channels 4.1 (Kir4.1) (Beverley & Pattnaik, 2022; A. Bringmann et al., 2006). Kir4.1 channels help stabilize the retinal membrane potential and manage K^+^-glutamate levels. (Connors & Kofuji, 2006; Katoozi et al., 2020; Li et al., 2021; Reichenbach & Bringmann, 2013). Diabetes has been shown to decrease Kir4.1 expression, leading to MC swelling and altered Kir4.1 distribution (Luo et al., 2019), which compromises MC function and disrupts retinal physiology (Lassiale et al., 2016). Such dysregulation in Kir4.1 can increase neuronal hyperexcitability (Amaratunga et al., 1996; Nwaobi et al., 2016) and impair K^+^ buffering (A. Bringmann et al., 2006; Kofuji et al., 2002). In AD, reduced Kir4.1 expression has been observed in postmortem brain samples with amyloid accumulation and mouse models of AD, suggesting a link between Kir4.1 dysfunction and AD pathology (Wilcock et al., 2009).

MCs are vital for maintaining retinal function, and their dysfunction, driven by factors like *APOE4*, can significantly impact retinal health and contribute to disease progression. However, this hypothesis has yet to be examined in the context of AD and *APOE4*. In this study, we have used *APOE4*-knock in (KI) mice and rat MC line (rMC-1) expressing *APOE4* to investigate the effects of *APOE4* on MCs and Kir4.1 channels. We aim to understand how *APOE4* influences mitochondrial dynamics and other cellular functions critical to retinal and neural health.

## 2 METHODS

### 2.1 Animals

The humanized *APOE*-KI mice were created via gene targeting, in which the native mouse *Apoe* gene was replaced with the human *APOE3* or *APOE4* gene. These mice were developed by the Model Organism Development and Evaluation for Late-Onset Alzheimer’s Disease (MODEL-AD) consortium. These mice were homozygous for either the *APOE3* (*3/3*) or *APOE4* (*4/4*) alleles. Hereafter, we will refer to them as *APOE3* mice and *APOE4* mice, respectively. The mice were housed at the animal care facility of the Eugene and Marilyn Glick Eye Institute, Indiana University, Indianapolis, IN, USA. All the animals were maintained under standard physiological conditions, including a 12-hour light/dark cycle, with continuous access to food and water. All experiments followed the Guiding Principles in the Care and Use of Animals (National Institute of Health) and the Association for Research in Vision and Ophthalmology (ARVO) Statement for the Use of Animals in Ophthalmic and Vision Research. Experiments were conducted on the animals aged between 52-57 weeks of age.

### 2.2 Whole-cell voltage-clamp recording

At 52-57 weeks of age, mice were euthanized, and after the eyes were enucleated, the retinas were isolated. The retinas were then incubated in Ringer’s solution containing 0.3mg/ml papain and 2.5mM L-cysteine for 30 minutes at 37^0^C. Following this, the retinas were briefly incubated at Dulbecco’s Modified Eagle’s Medium (DMEM, Thermo Fisher Scientific, MA, USA) with 10% fetal bovine serum (FBS, Thermo Fisher Scientific, MA, USA) and 0.2mg/ml DNase-1 at room temperature (RT), and the tissue was gently triturated. The resulting cell suspension was layered over a discontinuous Percoll gradient (10%, 20%, 30%, and 50% Percoll) and centrifuged at 800g for 5 minutes. The fraction enriched in MCs, found at the top of the 30% Percoll layer, was collected, washed with DMEM containing 10% FBS, and transferred to Poly-L-Lysine and laminin-coated coverslips to promote cell adhesion. Whole-cell patch clamp recordings were then performed in the voltage-clamp mode to measure Kir4.1 current in the MCs, as previously described (Thompson et al., 2018).

### 2.3 Cell Culture and Transfections

rMC-1 was generously provided by Dr. Vijay Sarthy, Northwestern University, Chicago, IL, USA. The cells were cultured in low glucose, no phenol red, DMEM (Thermo Fisher Scientific, MA, USA) supplemented with 10% FBS, 1% L-glutamine (Corning, VA, USA) and 1% antibiotic-antimycotic (Thermo Fisher Scientific, MA, USA). rMC-1 were grown in DMEM overnight and transfected with 1µg of plasmids encoding human *APOE* isoforms: pCMV4-*APOE2* (Cat. #87085, addgene, MA, USA), pCMV4-*APOE3* (Cat. #87086, addgene), and pCMV4-*APOE4* (Cat. #87087, addgene). Cells transfected with empty vector (EV, pCMV4-HA, Cat. #27553, addgene) were used as a control. The human APOE plasmids do not have any tag, while the EV has HA tag. Transfections were performed using Lipofectamine 3000 (L3000-008, Invitrogen, Thermo Fisher Scientific, MA, USA) following the manufacturer’s protocol. Cells were collected 24 hours post-transfection for mRNA, protein, and flow cytometry analyses. The validation of transfection was performed using immunofluorescence staining.

### 2.4 Immunofluorescence

At 52-57 weeks of age, mice were euthanized, and their eyes were fixed in 4% Paraformaldehyde (PFA) solution for 15 minutes at RT, followed by rinsing with phosphate-buffered saline (PBS). The retinas were then isolated from the fixed eyes, embedded in 3% agarose, and sectioned with a vibratome. Agarose sections were washed in a buffer containing 3% Dimethyl Sulfoxide (DMSO, Thermo Fisher Scientific, MA, USA) and 0.3% TritonX-100 (Thermo Fisher Scientific, MA, USA) in PBS, then blocked for 2 hours at RT with 5% goat serum diluted in washing buffer. Sections were then incubated overnight at 4^0^C with primary antibodies, including Kir4.1 (Cat. #APC-035-GP, Alomone Labs, 1:200), glutamine synthetase (GS, Cat. #MAB302, Millipore, 1:200) and TOMM20 (Cat. #MA5-32148, Invitrogen, 1:100). Next day sections were incubated with appropriate secondary antibodies. To validate transfections, rMC-1 was seeded on an 8-well chamber slide and transfected as described earlier. Later, cells were fixed with 4% PFA for 15 minutes, permeabilized using 0.3% TritonX-100 diluted in PBS, and blocked for 1 hour with 5% goat serum diluted in permeabilization solution. The cells were then incubated O/N at 4^0^C with Anti-HA (Cat. #26183, Invitrogen, 1:200), APOE (Cat. #ab52607, Abcam, 1:100), APOE3 (Cat. #MAB41442-SP, Novus Biologicals, CO, USA, 1:100) and APOE4 (Cat. #NBP1-49529SS, Novus Biologicals, 1:100) antibody, followed by washing and a 2-hour incubation with appropriate secondary antibody next day. Transfected rMC-1 were stained with TOMM20 to check the effect of *APOE* isoforms on mitochondria. Images were captured using a Zeiss LSM-700 confocal microscope (Carl Zeiss MicroImaging, Germany). The fluorescence intensities from the retinal sections for Kir4.1, TOMM20, and GS were calculated by subtracting fluorescence intensity from the secondary antibody control. The integrated density per cell area for TOMM20 staining in rMC-1 was calculated from total Z-stack projections using Fiji ImageJ software.

### 2.5 qRT-PCR for mRNA Analysis

Total RNA was extracted using Trizol reagent (Thermo Fisher Scientific, MA, USA) following manufacturer’s instructions, and 1μg of RNA was then reverse-transcribed with the SuperScript VILO cDNA synthesis kit (Thermo Fisher Scientific, MA, USA). Quantitative real-time PCR was performed using gene-specific primers, TaqMan Fast Universal Master Mix (Thermo Fisher Scientific, MA, USA), and the Viia7 Real-Time PCR system (Thermo Fisher Scientific, MA, USA) to measure mRNA levels. mRNA expression levels for each gene were normalized to the housekeeping gene *Bact* (Rn00667869_m1). Primers used were *Kcnj10* (gene for *Kir4.1*; Rn00581058_m1), *Mfn1* (gene for Mitofusin-1, Rn00594496_m1), *Mfn2* (gene for Mitofusin-2, Rn00500120_m1), and *Dnm1* (gene Dynamin-1, Rn00586466_m1).

### 2.6 Western Blotting

RIPA buffer (#R0278, Sigma-Aldrich Corp.) containing a protease inhibitor mixture was used to lyse rMC-1. Protein concentrations were measured using BCA assay (Pierce, Thermo Fisher Scientific), and equal amounts of protein were loaded onto a 4-12% Bis-Tris gel (Novex, Thermo Fisher Scientific) for separation. Proteins were then transferred onto PVDF membrane and blocked with 4% BSA in TBST buffer. The membranes were probed with primary antibodies against α-Tubulin (Cat. #T9026, 1:2000; Sigma-Aldrich Corp.) and Kir4.1 (Cat. # APC-035, 1:2000, Alomone Labs) O/N at 4^0^C. The next day, membranes were incubated with secondary peroxidase antibodies at RT for 2 hours. Bands were visualized using an ECL2 western blotting substrate (Thermo Fisher Scientific) and scanned with a Typhoon FLA 9500 laser scanner (GE Healthcare Life Sciences, PA, USA). Protein band intensities were quantified using ImageJ software. Integrated optical density (IOD) was calculated by taking ratio of Kir4.1 and α-tubulin.

### 2.7 Mitochondrial membrane potential (ΔΨm)

24 hours after transfections rMC-1 were resuspended in 1ml of DMEM at ∼1×10^6^cells/ml and incubated with 2μM JC-1 (5’,6,6’-tetrachloro-1,1’,3,3’-tetraethylbenzimidazolylcarbocyanineiodide, Molecular Probes, Invitrogen, CA, USA) for 30 minutes at 37^0^C in the dark, following the manufacturer’s instructions. Unstained and EV-treated cells were used as a control. For each sample, 100,000 gated events were acquired using a BD LSR Fortessa^TM^ cell analyzer (BD Biosciences, San Jose, CA) with 582/15nm (PE) filters for JC-1 aggregates and 525/50nm (FITC) filters for JC-1 monomers. Data were analyzed using FlowJo software (TreeStar, OR, USA). Dead cells and debris were excluded based on forward and side scatter, and all analyses were gated on unstained cells based on forward and side scatter morphology.

### 2.8 Mitochondrial reactive oxygen species (ROS) measurement

rMC-1 were incubated with 1μM MitoSox Red (MSR) mitochondrial superoxide indicator (Molecular Probes, Invitrogen, CA, USA) for 30 minutes at 37^0^C in the dark, according to the manufacturer’s instructions. Unstained and EV-transfected cells were used as controls. For each sample, 100,000 gated events were recorded on a BD LSRFortessa™ (BD Biosciences, NJ, USA) cell analyzer using a 610/20nm (PE-Texas Red) filter. Data analysis was conducted using FlowJo™ v10.10 software (BD Life Sciences, NJ, USA). Dead cells and debris were excluded based on forward and side scatter, and analyses were gated on unstained cells.

### 2.9 Mitoquinone mesylate (MitoQ) treatment

1µM MitoQ (Cat. #317102, MedKoo Biosciences Inc., NC, USA) was prepared according to the manufacturer’s instructions. To find the optimal dose of MitoQ, we made three concentrations of MitoQ: 0.5µM, 1µM, and 2µM. 24 hours after transfections, rMC-1 were washed twice with PBS and incubated in serum free medium (SFM) for 2 hours before MitoQ treatments. A 1:1 ethanol-to-water mixture was used as a vehicle. Cells were then treated with different concentrations of MitoQ and vehicle and incubated for 24 hours at 37^0^C. After 24 hours of treatment, gene expression of Kir4.1 was measured using qRT-PCR as described previously. All the remaining experiments, such as western blot and MSR flow cytometry, were performed with 1µM of MitoQ as described earlier.

### 2.10 Alamar Blue Viability Assay

rMC-1 were seeded in a flat, clear bottom 96 well plate at a density of 25,000 cells/well in 100µl DMEM, and transfections were carried out as mentioned previously. The following day, transfected cells were treated with MitoQ and vehicle, while non-transfected cells received a 20% DMSO treatment as a positive control. After 24 hours of MitoQ treatment, the medium was replaced with 100µl SFM, and 11.1µl of Alamar Blue (Bio-Rad, CA, USA) was added to each well. Cells were incubated with Alamar blue for 4 hours at 37^0^C. Fluorescence was measured using a Synergy H1 plate reader (BioTek, Winooski, VT) with an excitation wavelength of 560nm and an emission wavelength of 590nm. Raw fluorescence values were normalized to the fluorescence of DMSO-treated control cells. The % viable cells was calculated by taking a ratio of MitoQ-treated cells and vehicle-treated cells.

### 2.11 Statistical Analysis

For the animal studies, we used n= 9 *APOE3* mice (26 cells) and n= 8 *APOE4* mice (33 cells) for whole-cell voltage-clamp recording and n=3 animals per group for immunofluorescence staining. For *in vitro* experiments n= 3-5 independent experiments were performed with 3 technical replicates per experiment. Data were expressed as mean± standard error of the mean (SEM) and analyzed with GraphPad Prism 10.0.1 for Windows (San Diego, California; www.graphpad.com). t-test was used to compare fluorescence intensities and current densities for whole-cell voltage clamp recording. Intergroup companions were conducted using one-way ANOVA followed by Tukey’s multiple comparison test. A p-value of less than 0.05 was considered statistically significant. *p<0.05, **p<0.01, ***p<0.001, and ****p<0.0001.

## 3 RESULTS

### 3.1 *APOE4* allele causes structural and functional deficits in the MCs

To study the effect of *APOE4* on MCs, agarose-embedded retinal sections were stained for Kir4.1 and GS (Figure 1a). Kir4.1 is localized to the vitreal border and in perivascular processes in the outer retina (Connors & Kofuji, 2006). *APOE4* retinas did not show Kir4.1 in vitreal border compared to the *APOE3* retinas. There was a marked reduction in both Kir4.1 (p= 0.0019) and GS (p< 0.0001) in *APOE4* retinas compared to the *APOE3* retinas (Figure 1b), suggesting impaired structural integrity in MCs is associated with the *APOE4* allele. To further assess Kir4.1 function, we conducted whole-cell patch clamp recordings in the voltage-clamp mode. These recordings showed a significant reduction (∼1.6 fold) in Kir4.1 current density in *APOE4* compared to *APOE3* MCs, indicating compromised K^+^ buffering ability in *APOE4* MCs (p= 0.0001, Figure 1c).

**Figure 1:**
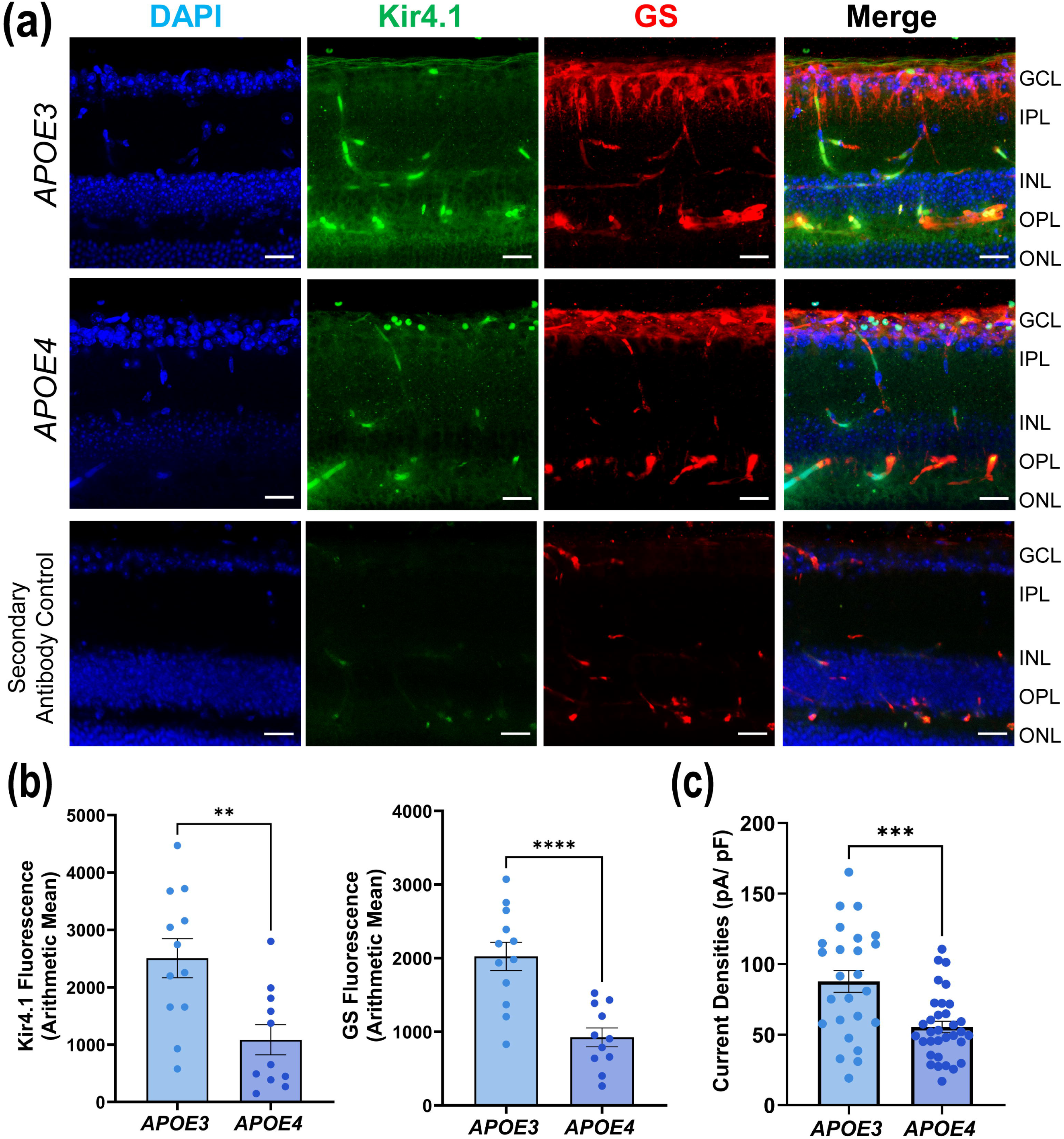
*APOE4* causes deficits in the Kir4.1. (a) Representative images of retinal slices showing Glutamine synthase (GS) and Kir4.1 staining pattern in *APOE3* and *APOE4* mice, scale 20µm (n: *APOE3*= 3, *APOE4*= 3). (b) Bar graph showing quantification of immunofluorescence for Kir4.1 and GS (n: 11-12 images/ group). (c) Current densities of Kir4.1 from freshly isolated Müller cells from *APOE3* and *APOE4* mice. Currents were elicited by a 50-ms hyperpolarization of −140 mV from a holding potential of −60 mV. (n: *APOE3*= 26 cells/ 9 mice, *APOE4*= 33 cells/ 8 mice). Values are expressed as mean ± SEM. Unpaired t-test was used for statistical analysis. **p<0.01, ***p<0.001, ****p<0.0001.

### 3.2 *APOE4* allele leads to mitochondrial dysfunction

Mitochondrial impairment is well-documented in AD, and Figure 2a illustrates a proposed hypothesis by which *APOE4* may drive induced MC dysfunction via mitochondrial impairment. The *APOE4* allele disrupts mitochondrial gene expression by acting as a transcriptional factor or by directly interacting with mitochondria, altering metabolism and fusion/fission balance, resulting in reduced ΔΨm, ROS generation, and subsequent mitochondrial dysfunction (Chang et al., 2005). These mitochondrial deficits may result in the downregulation of Kir4.1 channels, compromising MC function. This pathway suggests a potential mechanism by which *APOE4* contributes to inflammation, accelerated aging, and cell death, linking to cellular degeneration in AD. To investigate whether *APOE4* also induces mitochondrial deficits in the retina, we stained retinal sections from *APOE3* and *APOE4* mice with TOMM20 (a mitochondrial marker) (Figure 2b). Retinas from *APOE4* mice displayed a marked reduction in TOMM20 fluorescence (p= 0.0404) along with GS (p= 0.0005) compared to *APOE3* retinas (Figure 2c), indicating a decrease in mitochondrial content or function. These findings suggest that the *APOE4* allele may contribute to the mitochondrial dysfunction in retinal cells, potentially linking broader cellular impairments seen in AD.

**Figure 2:**
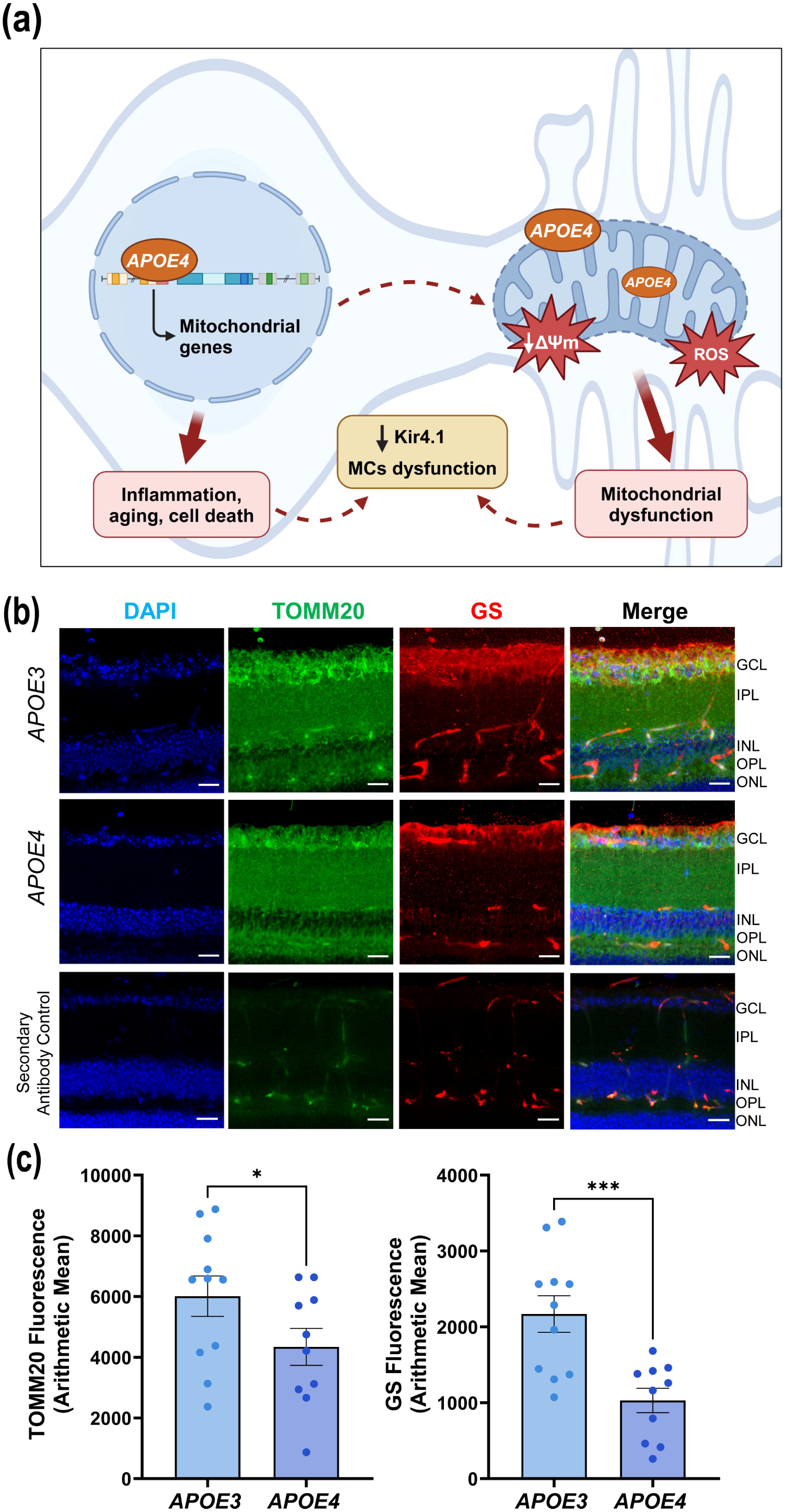
Mitochondrial dysfunction in *APOE4*. (a) An illustration suggests a potential mechanism by which *APOE4* disrupts Kir4.1 expression and causes Müller cell dysfunction. *APOE4* may disrupt nuclear-encoded mitochondrial gene expression, reducing membrane potential (ΔΨm) and increasing reactive oxygen species (ROS). These changes can impair mitochondrial function, resulting in the downregulation of Kir4.1 channels in Müller cells and driving inflammation, aging, and cell death, contributing to retinal dysfunction in Alzheimer’s disease. (b) Representative images of retinal slices showing Glutamine synthase (GS) and TOMM20 staining pattern in *APOE3* and *APOE4* mice, scale 20µm (n: *APOE3*= 3, *APOE4*= 3). (c) Bar graph showing quantification of immunofluorescence for TOMM20 and GS (n: 10-11 images/ group). Values are expressed as mean ± SEM. Unpaired t-test was used for statistical analysis. *p<0.05, ***p<0.001.

### 3.3 *APOE4* transfected rMC-1 have lower Kir4.1 gene and protein expression

To further confirm our findings from the mouse model, we created an *in vitro* model by transfecting rMC-1 with *APOE2*, *APOE3*, or *APOE4*, using an EV as a control (Figure 3a). While humans have three APOE variants, rats have only one, which contains arginine at 112 (https://web.expasy.org/variant_pages/VAR_000652.html), unlike human *APOE4* and rat APOE is similar to human *APOE3*. First, we validated the transfection using immunofluorescence (Supplementary Figure 1). The staining showed that rMC-1 transfected with *APOE2*, *APOE3*, or *APOE4* plasmids exhibited distinct intracellular staining corresponding to the expressed APOE proteins, while the staining for anti-HA showed transfection with EV, confirming the efficacy of the transfection and the expression of human APOE isoforms in rMC-1. In line with the observations in retinal tissue, rMC-1 transfected with *APOE4* showed a significant decrease in Kir4.1 gene expression (Figure 3b) compared to the cells transfected with EV (p= 0.0124) or *APOE2* (p= 0.0123) or *APOE3* (p= 0.0363). Western blot analysis supported these results, revealing a marked reduction in Kir4.1 protein levels in *APOE4*-transfected rMC-1 (Figure 3c) compared to the EV (p= 0.0032) or *APOE2* (p= 0.0020) or *APOE3* (p= 0.0340).

**Figure 3:**
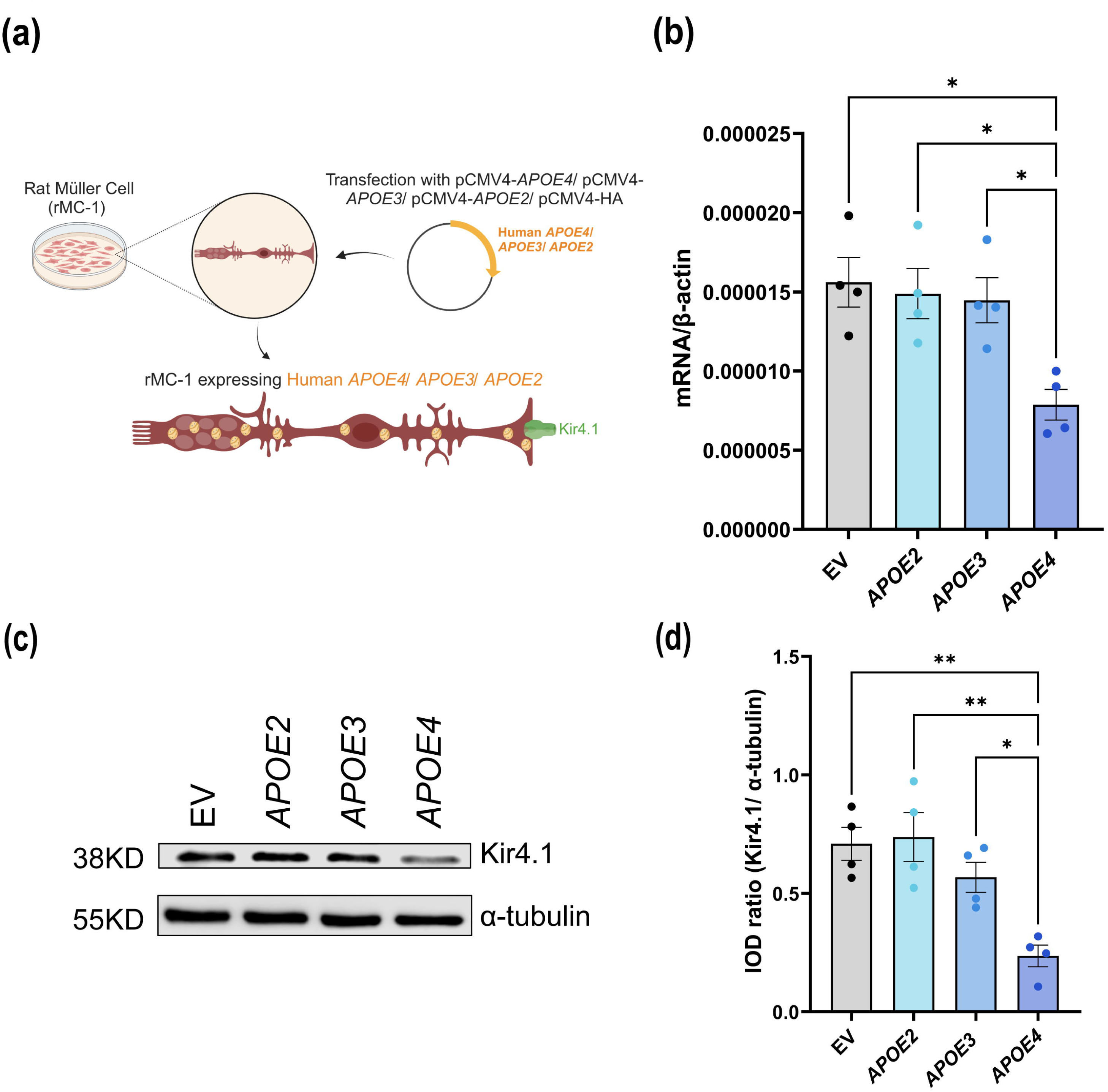
*APOE4* decreases Kir4.1 gene and protein expression in rMC-1. (a) Schematic showing generation of rMC-1 expressing human *APOE* isoforms. rMC-1 were transiently transfected with human *APOE2*/ *APOE3*/ *APOE4* and EV was used as a control. (b) mRNA expression of *Kcnj10* gene for Kir4.1 normalized to housekeeping gene β-actin. (c) Representative western blots of Kir4.1 expression and quantification of integrated optical density (IOD) ratio of Kir4.1 and α-tubulin showing decreased protein expression of Kir4.1 in *APOE4*-transfected rMC-1. Values are expressed as mean ± SEM. One-way ANOVA followed by Tukey’s multiple comparison test was used for statistical analysis. *p<0.05, **p<0.01. (n: 4 independent experiments)

### 3.4 Mitochondrial deficits in *APOE4* transfected rMC-1

We sought to assess mitochondrial health in *APOE4* transfected rMC-1. We stained rMC-1 transfected with EV/ *APOE2/ APOE3/ APOE4* with TOMM20 (Figure 4a) and found that TOMM20 staining intensity (Figure 4b) is decreased in rMC-1 transfected with *APOE4*, further confirming findings from *in vivo* staining. The TOMM20 staining intensity was found to be significantly decreased in *APOE4* transfected rMC-1 compared to EV/ *APOE2*/ *APOE3* (p= <0.0001). We examined mRNA expression of mitochondrial fusion genes *Mfn1* and *Mfn2* and fission gene *Dnm1* (Figure 4b). Results showed that *APOE4* transfection led to significant downregulation of *Mfn1*, *Mfn2* and *Dnm1* expression compared to EV (p= 0.0097, 0.0002, 0.0022) or *APOE2* (p= 0.0749, 0.0078, 0.0924) or *APOE3* (p= 0.0313, 0.0253, 0.0398). These findings collectively reinforce the role of *APOE4* in mitochondrial dysfunction and Kir4.1 downregulation, suggesting a consistent mechanism of MC dysfunction both *in vivo* and *in vitro*.

**Figure 4:**
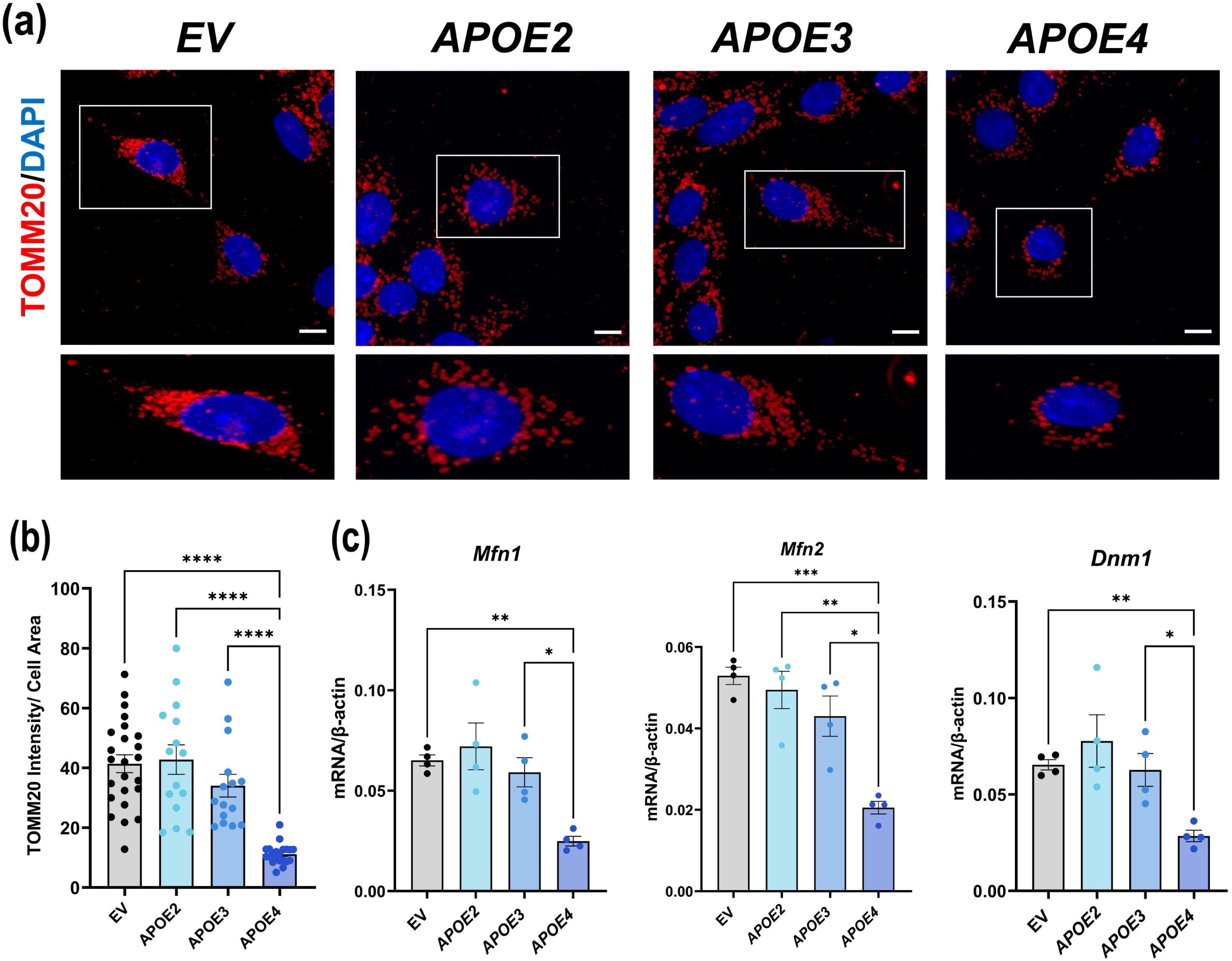
*APOE4* decreases mitochondrial gene expression in rMC-1. (a) Representative images of rMC-1 transfected with human *APOE2*/ *APOE3*/ *APOE4*/ EV showing decreased TOMM20 staining pattern in *APOE4*-transfected rMC-1, scale: 20µm. (n: 3 independent experiments) (b) Quantification of TOMM20 staining intensity per cell area. (n: 15-24 cells/ condition) (c) mRNA expression of *Mfn1*, *Mfn2* and *Dnm1*, showing *APOE4*-transfected rMC-1 reduced *Mfn1*, *Mfn2* and *Dnm1* gene expression as compared to EV/ *APOE2*/ *APOE3* transfected rMC-1. (n: 4 independent experiments) Values are expressed as mean ± SEM. One-way ANOVA followed by Tukey’s multiple comparison test was used for statistical analysis. *p<0.05, ****p<0.0001.

### 3.5 *APOE4* impairs mitochondrial membrane potential (ΔΨm) in rMC-1

To assess ΔΨm in rMC-1 expressing different *APOE* isoforms, we performed JC-1 flow cytometry analysis. The results showed a notable decrease in ΔΨm in cells transfected with *APOE4* compared to those transfected with EV, *APOE2*, or *APOE3* (Figure 5a), suggesting that *APOE4* negatively impacts mitochondrial function. Quantitative analysis revealed a significant reduction in the ratio of red (aggregated JC-1, indicating normal ΔΨm) to green (monomeric JC-1, indicative of mitochondrial depolarization) fluorescence in *APOE4*-expressing rMC-1 compared to EV (p= <0.0001) or *APOE2* (p= <0.0001) or *APOE3* (p= 0.0001) (Figure 5b). This shift towards green fluorescence in *APOE4* transfected cells highlights a loss of mitochondrial membrane potential, a hallmark of mitochondrial dysfunction.

**Figure 5:**
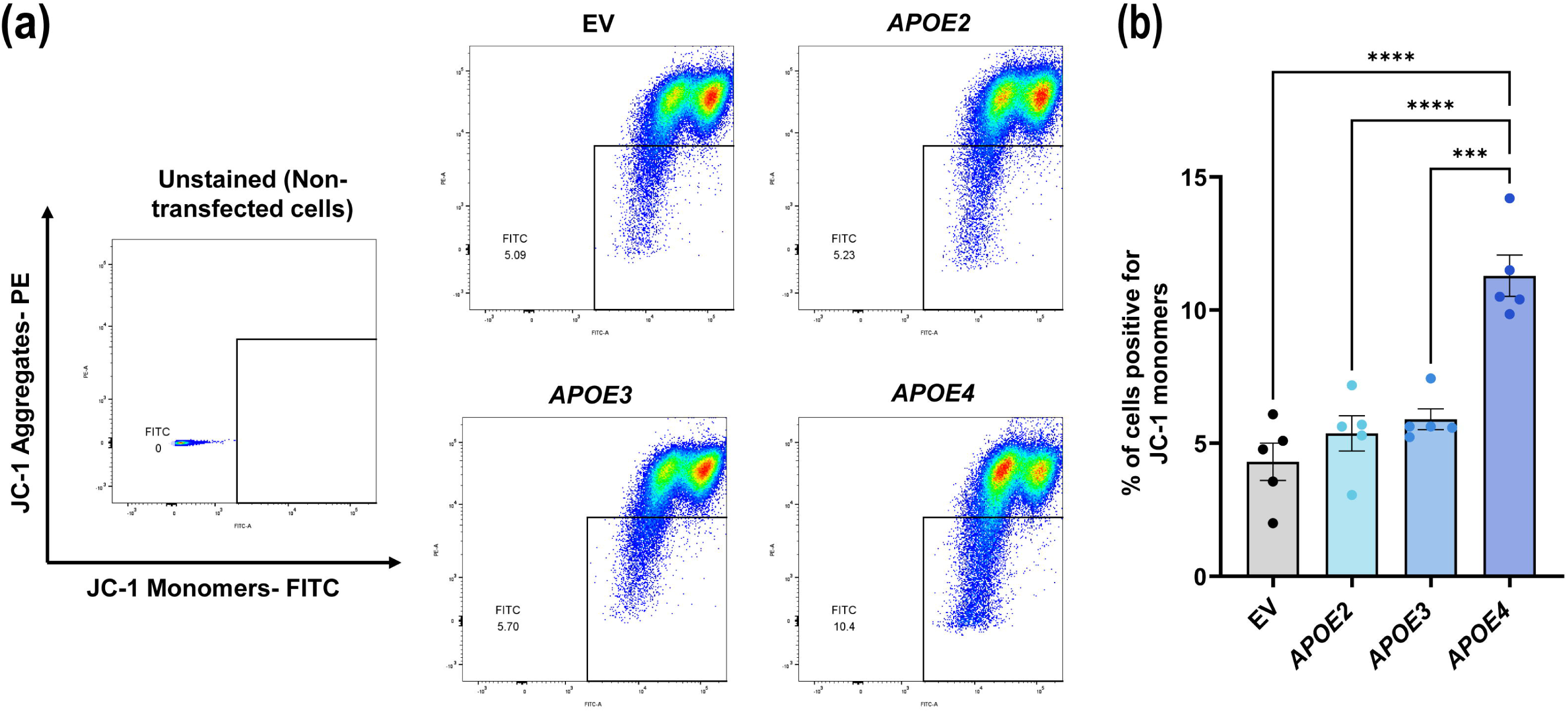
*APOE4* decreases Mitochondrial membrane potential (ΔΨm) in rMC-1. (a) Representative images of unstained rMC-1 and rMC-1 transfected with EV/ *APOE2*/ *APOE3*/ *APOE4* and analyzed on a flow cytometer with 525/50nm and 582/15nm bandpass emission filters. (b) Bar graph showing quantification of % of the cells positive for JC-1 monomers. Values are expressed as mean ± SEM (n: 5 independent experiments). One-way ANOVA followed by Tukey’s multiple comparison test was used for statistical analysis. ***p<0.001, ****p<0.0001.

### 3.6 *APOE4* increases mitochondrial ROS accumulation

To investigate oxidative stress within the mitochondria, we measured mitochondrial ROS levels in rMC-1 cells transfected with *APOE*4, using MSR flow cytometry analysis (Figure 6a). *APOE*4-expressing cells exhibited a significant increase in mitochondrial ROS production compared to cells expressing EV (p= 0.0003) or *APOE2* (p= 0.0032) or *APOE3* (p= 0.0282) (Figure 6b), indicating heightened oxidative stress, specifically associated with the *APOE*4 isoform. This elevation in ROS further underscores the mitochondrial impairments linked to *APOE*4, contributing to cellular stress and potential damage within the retinal environment.

**Figure 6:**
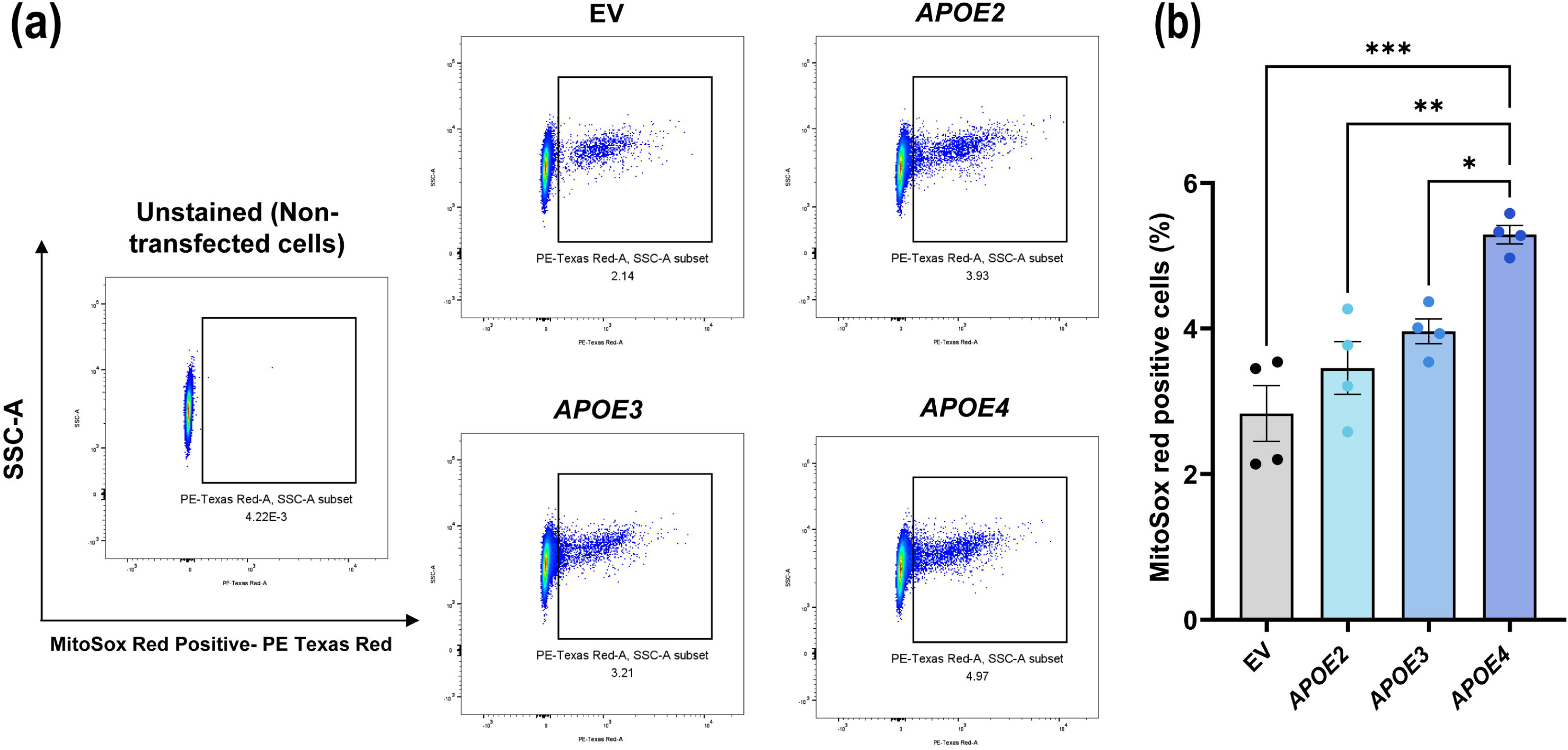
*APOE4* increases Mitochondrial ROS production in rMC-1. (a) Representative images of unstained rMC-1 and rMC-1 transfected with EV/ *APOE2*/ *APOE3*/ *APOE4* and analyzed on a flow cytometer with 610/20nm bandpass emission filter. (b) Bar graph showing quantification of % of MitoSox Red positive cells. Values are expressed as mean ± SEM (n: 4 independent experiments). One-way ANOVA followed by Tukey’s multiple comparison test was used for statistical analysis. *p<0.05, **p<0.01, ***p<0.001.

### 3.7 MitoQ treatment restores Kir4.1 expression in *APOE4*-transfected rMC-1

To investigate whether mitochondrial-targeted antioxidant MitoQ could mitigate the mitochondrial dysfunction and restore Kir4.1 expression in *APOE4*-transfected rMC-1, we treated these cells with MitoQ and assessed its impact on Kir4.1 levels. First, we tested 3 different concentrations of MitoQ 0.5µM, 1µM and 2µM and its effect on Kir4.1 gene expression. We found that 1µM MitoQ significantly increased *Kcnj10* mRNA expression in *APOE4*-transfected rMC-1 compared to the 0.5µM and 2µM concentrations (Supplementary Figure 2). Therefore, we performed the remaining experiments with 1µM MitoQ. Following 1µM MitoQ treatment, *APOE4*-transfected cells significantly improved *Kcnj10* gene expression (p= 0.0037, Figure 7a) compared to vehicle-treated *APOE4* cells. We observed *APOE4*-transfected cells have lower *Kcnj10* gene expression in vehicle-treated EV (p= 0.0069) or *APOE2* (p= 0.0463) or *APOE3* (p= 0.0239). Similarly, *APOE4*-transfected cells treated with MitoQ showed significantly improved Kir4.1 protein expression (p= 0.0041, Figure 7b) compared to vehicle-treated *APOE4* cells. This increase in Kir4.1 expression in MitoQ-treated *APOE4* cells brought them closer to levels observed in EV, *APOE2*, and *APOE3*-transfected cells. These findings suggest that MitoQ, by enhancing mitochondrial health, can partially rescue Kir4.1 expression in *APOE4*-expressing rMC-1.

**Figure 7:**
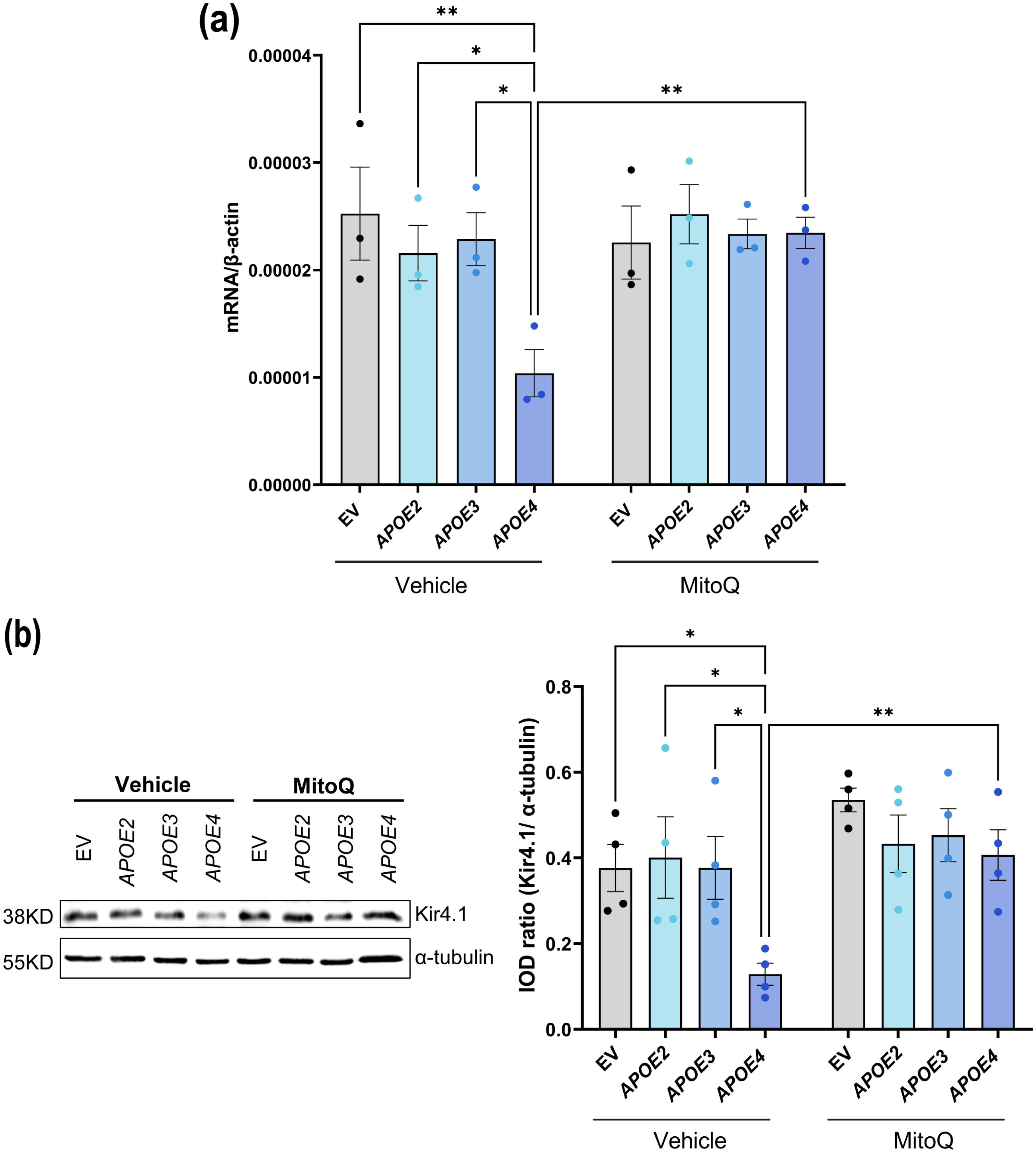
MitoQ restores Kir4.1 gene and protein expression in rMC-1 transfected with *APOE4*. (a) mRNA expression of *KCNJ10* gene for Kir4.1 normalized to housekeeping gene for β-actin after treating rMC-1 with 1µM MitoQ and vehicle. mRNA expression of Kir4.1 was significantly increased in *APOE4*-transfected rMC-1 upon treatment with 1µM MitoQ compared to the vehicle. (b) Representative western blots of Kir4.1 expression and quantification of IOD ratio of Kir4.1 and α-tubulin showing comparable protein expression of Kir4.1 in *APOE4*-transfected rMC-1 as compared to EV/ *APOE2*/ *APOE3* transfected rMC-1 after treating with 1µM MitoQ. Values are expressed as mean ± SEM. Two-way ANOVA followed by Tukey’s multiple comparison test was used for statistical analysis. *p<0.05, **p<0.01. (n: 3-4 independent experiments)

### 3.8 MitoQ treatment does not induce toxicity in *APOE4*-transfected rMC-1

To evaluate the safety and potential toxicity of MitoQ treatment on rMC-1, we performed a viability assay using Alamar Blue in cells transfected with EV, *APOE2*, *APOE3*, and *APOE4* (Supplementary Figure 3a). Following MitoQ treatment, we observed no reduction in cell viability compared to the vehicle-treated controls across all the groups. Each group including EV, *APOE2*, *APOE3*, and *APOE4*-transfected cells-demonstrated ∼100% viability after MitoQ exposure (Supplementary Figure 3b), indicating that the treatment does not induce cytotoxic effects at the applied concentrations. This confirms that MitoQ is well-tolerated by rMC-1 cells and suitable for further experiments to improve mitochondrial function and cellular health in *APOE4*-expressing cells.

### 3.9 MitoQ reduces mitochondrial ROS in *APOE4-*transfected rMC-1 to levels comparable with *APOE2*/ *APOE3-*transfected rMC-1

To further examine MitoQ’s impact on mitochondrial oxidative stress in *APOE4-* transfected cells, we conducted MSR flow cytometry to measure mitochondrial ROS levels following MitoQ treatment (Figure 8). Results indicated that MitoQ-treated *APOE4-*transfected rMC-1 exhibited a significant reduction in mitochondrial ROS compared to vehicle-treated *APOE4* rMC-1 (p= 0.0162, Figure 8a, b). Also, vehicle-treated *APOE4*-transfected rMC-1 has elevated ROS levels as compared to EV (p= 0.0088) or *APOE2* (p= 0.0266), and though not significant with *APOE3* (p= 0.0239), there is a trend, suggesting *APOE3*-transfected cells has lower mitochondrial ROS compared to *APOE4*-transfected cells. Notably, this reduction brought mitochondrial ROS levels in *APOE4* cells down to levels comparable with those observed in EV, *APOE2*, and *APOE3*-transfected cells treated with either vehicle or MitoQ. These findings suggest that MitoQ effectively mitigates the elevated oxidative stress associated with the *APOE4* isoform, restoring mitochondrial ROS levels to those typical of *APOE2* and *APOE3* expression.

**Figure 8:**
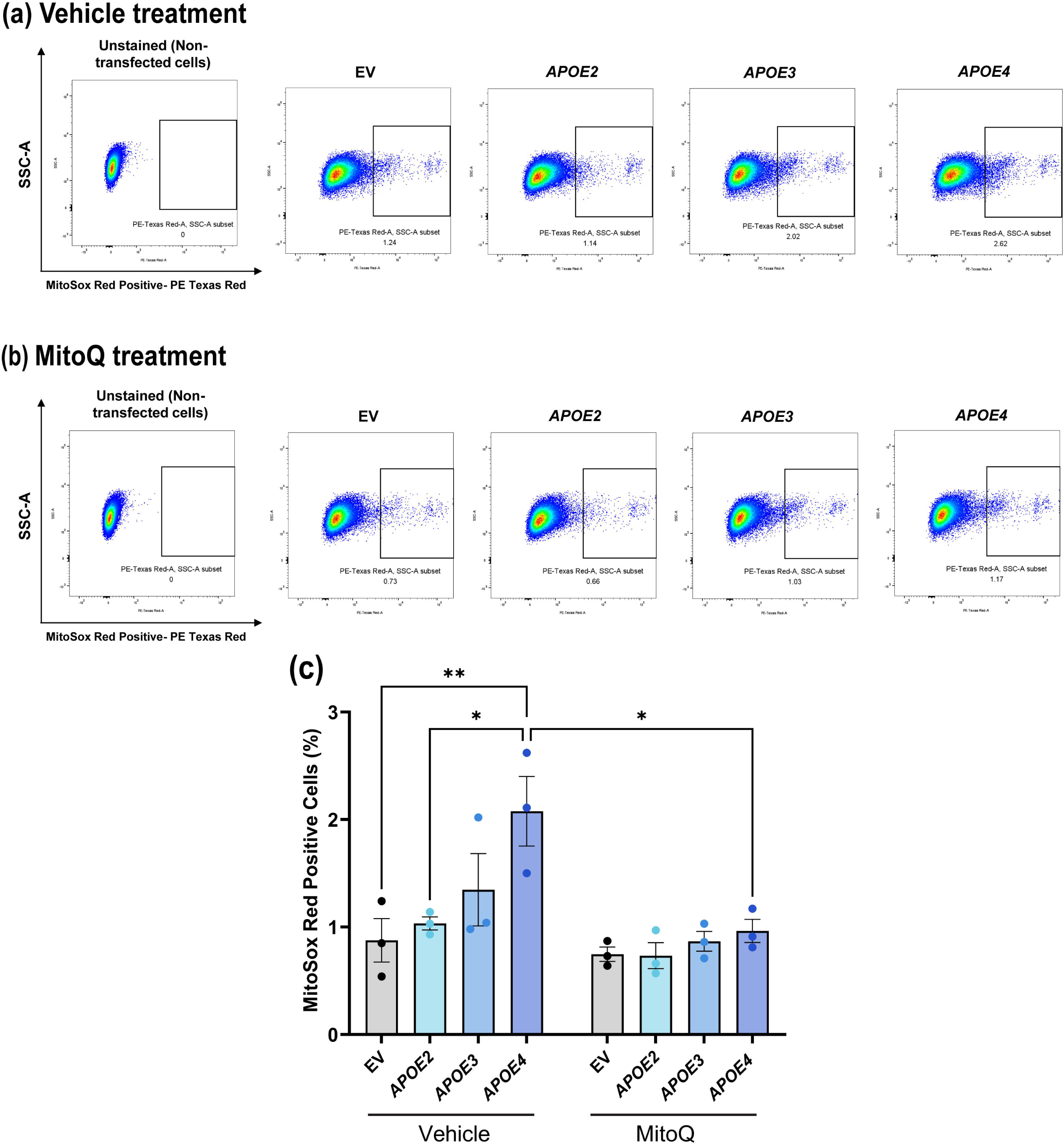
MitoQ decreases mitochondrial ROS in *APOE4-*transfected rMC-1. Representative images of unstained rMC-1 and rMC-1 transfected with EV/ *APOE2*/ *APOE3*/ *APOE4 and treate*d with (a) vehicle or (b) MitoQ (1µM). Cells were analyzed on a flow cytometer with 610/20nm bandpass emission filter. (c) Bar graph showing quantification of % of MitoSox Red positive cells. Mitochondrial reactive oxygen species (ROS) was decreased upon treating *APOE4-*transfected rMC-1 with 1µM MitoQ. Values are expressed as mean ± SEM (n: 3 independent experiments). One-way ANOVA followed by Tukey’s multiple comparison test was used for statistical analysis. *p<0.05, **p<0.01.

## 4 DISCUSSION

Our study shows that the *APOE4* allele causes significant structural and functional deficits in MCs. These deficits are associated with mitochondrial content, disrupted gene expression, and increased ROS production in *APOE4*-expressing MCs. Additionally, we emphasize that targeting mitochondrial impairments with antioxidants like MitoQ may offer a promising strategy for reducing the progression of retinal and neurodegenerative diseases.

We observe a marked reduction in Kir4.1 channels and GS expression in *APOE4*-*KI* retinas compared to *APOE3*-*KI*, indicating that *APOE4* disrupts MC structural integrity. Functionally, these disruptions are compounded by a significant decrease in Kir4.1 channel current density, reflecting impaired K^+^ buffering capacity—a critical function of MCs. These findings align with previous studies showing reduced Kir4.1 mRNA levels in the medial temporal lobe of AD patients and with severe amyloid angiopathy, as well as reduced Kir4.1 mRNA and protein in APPSwDI/NOS2^−/−^ and APPSwDI mice (Wilcock et al., 2009). Conversely, upregulated Kir4.1 expression was found in the human AD cortex (Smith et al., 2022) and in human AD middle temporal gyrus (Liu et al., 2024). Similarly, increased Kir4.1 mRNA and protein expression were observed in the dentate gyrus around amyloid plaques in APP/PS1 mice; however, K^+^ levels in the hippocampus and cortex remained unchanged (Huffels et al., 2022), suggesting that Kir4.1 function remained intact. Of note, our research is the first to show damage to the Kir4.1 structure and function in *APOE4*-KI. Additionally, no previous studies have reported findings in human or mouse AD retinas.

The link between *APOE4* and Kir4.1 dysfunction underscores the importance of glial cells in AD pathology, where glial dysfunction often precedes neuronal loss. Using an *in vitro* model of rMC-1, we confirmed the *APOE4-specific* downregulation of Kir4.1 at both the transcript and protein levels. The consistency between *in vivo* and *in vitro* findings reinforces the relevance of our model and highlights the specific impact of the *APOE4* isoform on MC dysfunction. Notably, *APOE2* and *APOE3* transfections did not replicate these deficits, further emphasizing the unique pathogenic role of *APOE4*.

Our findings highlight the role of mitochondria in *APOE4*-mediated dysfunction, evidenced by reduced TOMM20 expression and mitochondrial content in *APOE4* retinas. Human *APOE4* carriers show lower MFN1, MFN2, DNM1, and sirtuin-3 in the brain (Yin et al., 2020), suggesting compromised mitochondrial biogenesis and function. We observed that *APOE4*-transfected rMC-1 exhibited significant reductions in the expression of mitochondrial fusion and fission genes (Mfn11, Mfn2, and Dnm1), indicating disrupted mitochondrial dynamics. Furthermore, these cells demonstrated impaired ΔΨm and increased mitochondrial ROS levels, hallmark features of mitochondrial dysfunction. These data are consistent with previous reports linking *APOE4* to disrupted mitochondrial biogenesis, oxidative stress, and deficits in ATP production (Liang et al., 2021; Orr et al., 2019; Troutwine et al., 2022) in N2a cells as well as brain tissues, critical contributors to neurodegenerative processes in AD. *APOE4* is associated with reduced mitochondrial antioxidant defenses, increased mitochondrial superoxide production, and oxidative damage to lipids and proteins (Marottoli et al., 2021). For AD patients carrying *APOE4*, elevated hydroxyl radials in the blood [9] and decreased cerebral oxygen consumption (Robb et al., 2022) have been observed, and neurons expressing *APOE4* demonstrate reduced ATP production (Orr et al., 2019). These findings underscore the profound impact of *APOE4* on mitochondrial dysfunction, highlighting its potential role in exacerbating oxidative stress and energy deficits that contribute to neurodegenerative processes in AD.

Our study provides promising evidence for the therapeutic potential of MitoQ, a mitochondrial-targeted antioxidant, in mitigating *APOE4*-induced MC dysfunction. MitoQ effectively reduced mitochondrial ROS levels in *APOE4*-transfected cells, restoring them to levels observed in *APOE3* and *APOE2*-transfected cells. Additionally, MitoQ treatment rescued Kir4.1 gene and protein expression in *APOE4* cells, bringing them closer to baseline levels seen in *APOE3*-expressing cells. These results suggest that MitoQ alleviates oxidative stress and addresses the downstream consequences of mitochondrial dysfunction, thereby improving MC health and function.

The retinal findings in this study reflect broader pathological changes observed in the brain during AD, underscoring the retina’s usefulness as a non-invasive model for studying neurodegenerative diseases. Considering the role of MCs in maintaining the blood-retinal barrier and supporting neuronal health, targeting mitochondrial dysfunction in these cells may offer a dual benefit of preserving retinal and brain health. The therapeutic effects of MitoQ observed here support its potential as a candidate for further clinical investigation settings. A study on 3xTg-AD mice has shown that MitoQ inhibited cognitive decline in these mice (Young & Franklin, 2019) and it was also shown to improve retinal function, and reduce oxidative stress, inflammation and apoptosis in a retinal ischemia-reperfusion injury rat model (Tang et al., 2022). Therefore, studying the effect of MitoQ, particularly on individuals carrying the *APOE4* allele who are at heightened risk for AD, might help improve their cognitive abilities.

While our findings offer crucial insights into *APOE4*-induced MC dysfunction, several questions remain unanswered. For instance, the degree to which *APOE4*-induced mitochondrial dysfunction directly drives other retinal pathologies, such as neuronal degeneration, requires further investigation. Furthermore, although MitoQ demonstrated effectiveness *in vitro, in vivo* studies are essential to validate its therapeutic potential and establish optimal dosing regimens. Future research should also examine whether other mitochondrial-targeted therapies or combination treatments could synergistically address *APOE4*-associated retinal and neurodegenerative impairments.

In summary, our study identifies a novel mechanism by which *APOE4* impairs MC function through mitochondrial dysfunction, resulting in reduced Kir4.1 expression and K^+^ buffering capacity. MitoQ’s ability to alleviate these deficits highlights the potential of targeting mitochondrial health as a therapeutic strategy for *APOE4*-associated retinal and neurodegenerative diseases. These findings underscore the need to explore mitochondrial therapeutics in the context of *APOE4* AD.

## ACKNOWLEDGMENTS

We thank Dr. Evan Cornett, Dr. Jeffrey Elmendorf, and Dr. Amelia Linnemann from Indiana University School of Medicine for their valuable suggestions and guidance.

## Data availability

The data that support the findings of this study are available from the corresponding author upon request.

## AUTHORS’ CONTRIBUTIONS

SA: Writing - Original Draft, Writing - Review & Editing, Conceptualization, Software, Validation, Formal analysis, Investigation. YX: Software, Validation, Formal analysis, Investigation, Writing - Review & Editing. NM: Writing - Review & Editing. QL: Writing - Review & Editing, Validation, Formal analysis, Investigation. TRC, AO, and BL: Resources, Writing - Review & Editing. TWC: Conceptualization, Writing - Review & Editing. AB: Conceptualization, Resources, Writing - Review & Editing, Supervision, Project administration, Funding acquisition.

## Abbreviations

*APOE2*: Apolipoprotein *E2*
*APOE3*: Apolipoprotein *E3*
*APOE4*: Apolipoprotein *E4*
MC: Müller cell
rMC-1: Rat Müller cell-1
Kir4.1: Inwardly rectifying K^+^ channels 4.1
ΔΨm: Mitochondrial membrane potential
MitoQ: Mitoquinone mesylate
EV: Empty vector

---

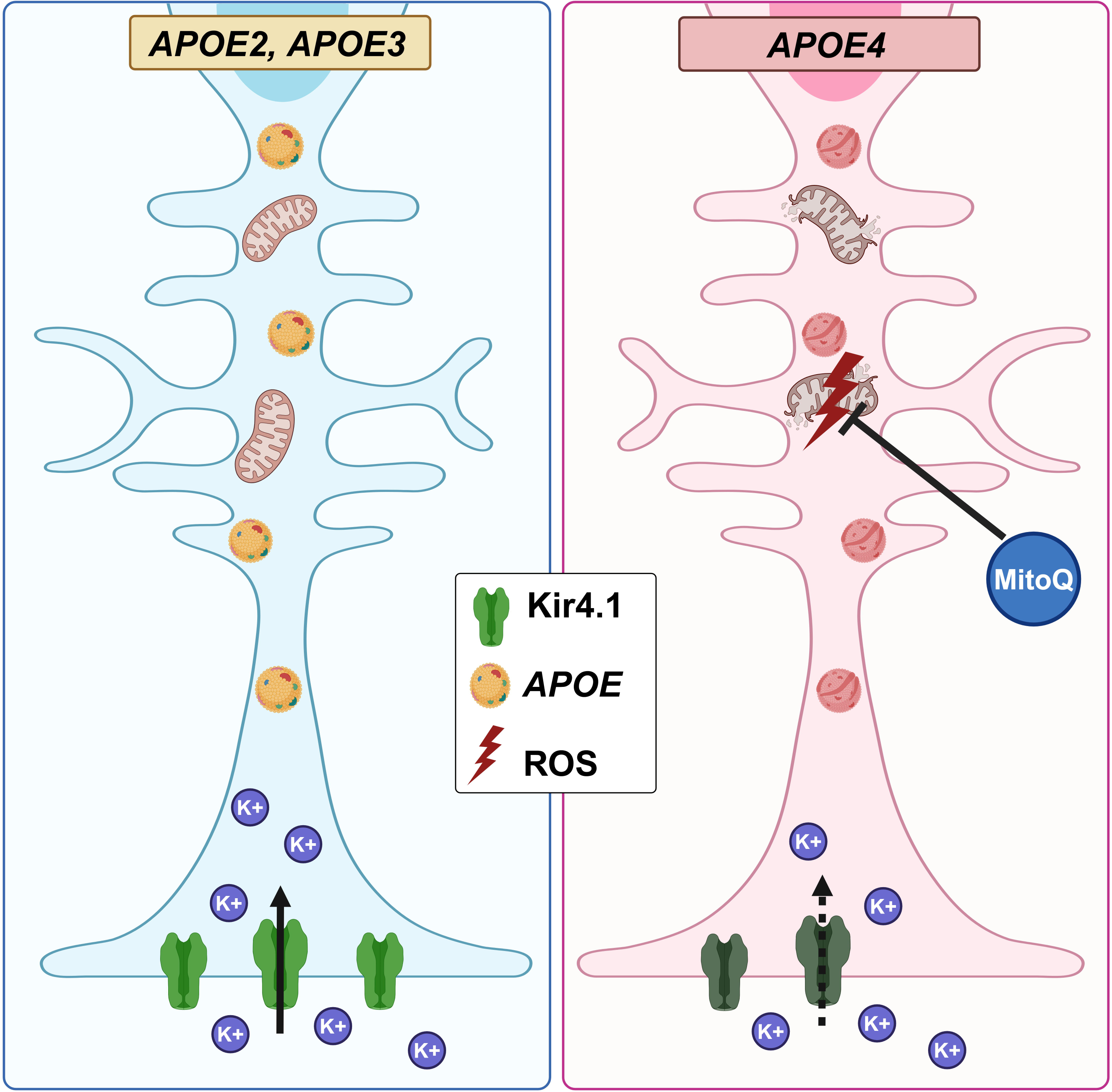

