## Supplementary material for "Müller glial Kir4.1 channel Dysfunction in *APOE4*-KI model of Alzheimer’s disease": Supplmentary Figure

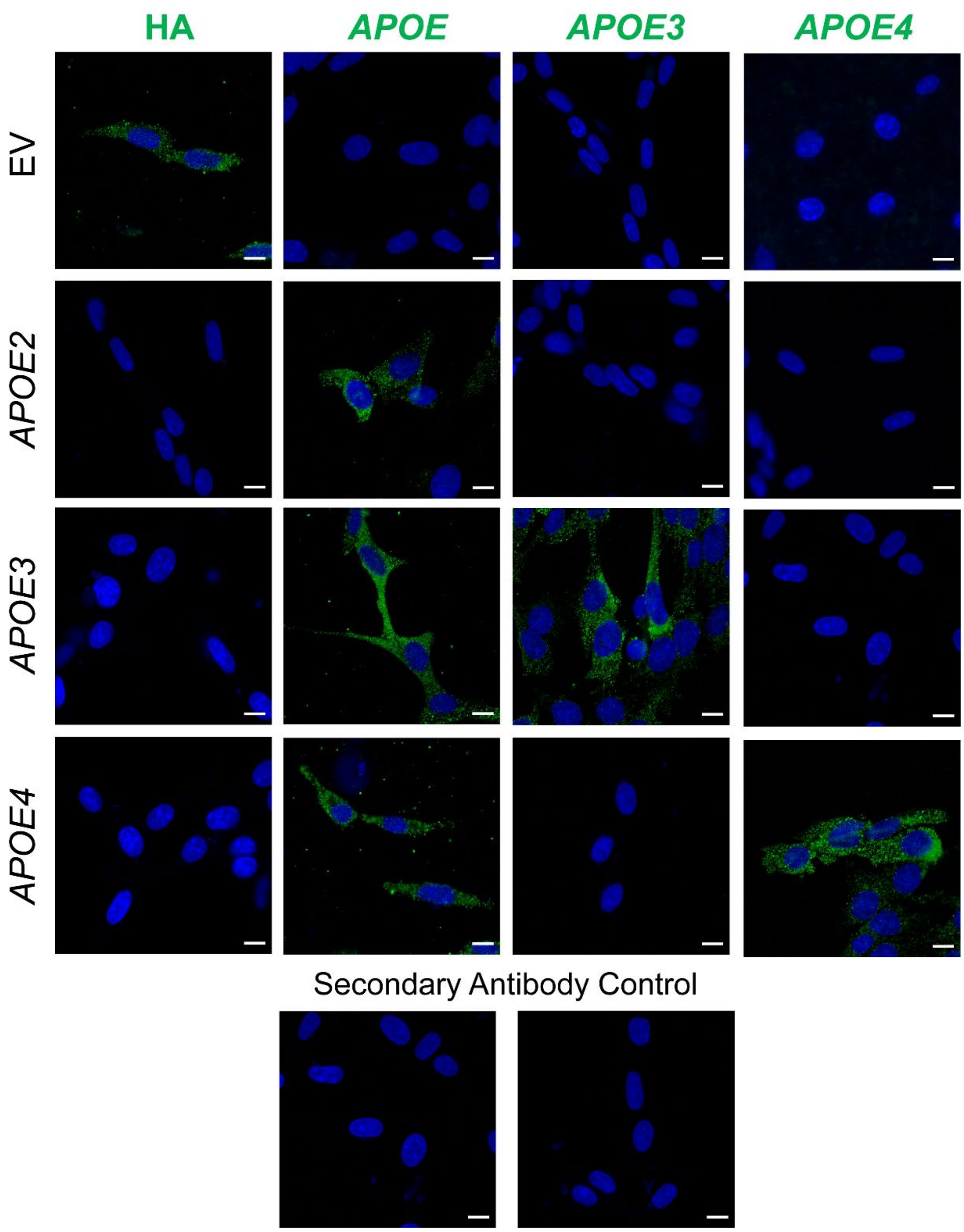


**Supplemental Figure 1: Confirmation of transfections.** Representative images of rMC-1 showing validation of transfection. rMC-1 transfected with EV or human *APOE2*/ *APOE3*/ *APOE4* were stained for each antibody: anti-HA (for EV), total APOE, APOE3 and APOE4. Scale 20μm. (n: 3 independent experiments)

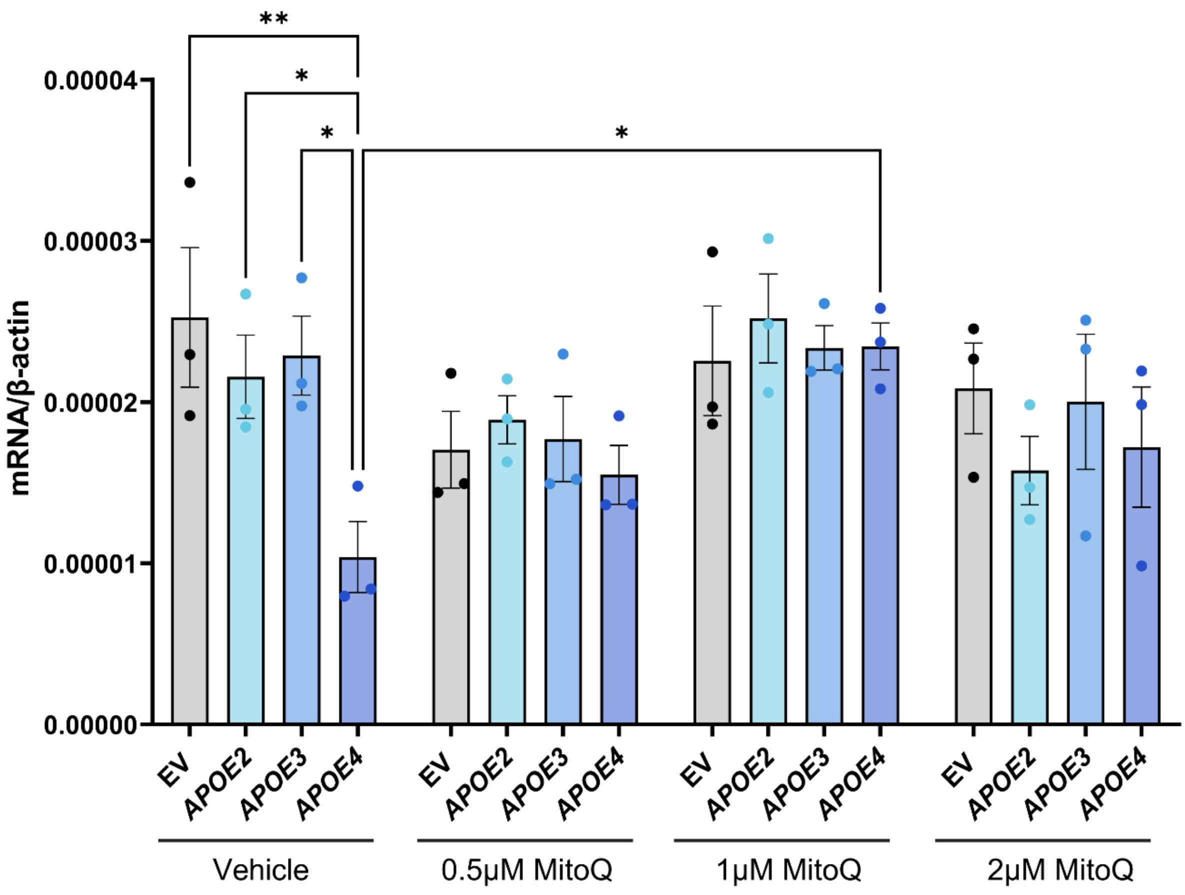


**Supplemental Figure 2: MitoQ (1μM) is the optimal dose to restore Kir4.1 gene expression in *APOE4*-transfected rMC-1.** mRNA expression of *KCNJ10* gene for *Kcnj10* normalized to housekeeping gene β-actin after treating rMC-1 with three different doses of MitoQ: 0.5 μM, 1μM and 2μM and vehicle. mRNA expression of Kir4.1 was significantly increased in *APOE4*-transfected rMC-1 upon treatment with 1μM MitoQ compared to the vehicle. Values are expressed as mean ± SEM. One-way ANOVA followed by Tukey's multiple comparison test was used for statistical analysis. *p<0.05, **p<0.01. (n: 3 independent experiments)

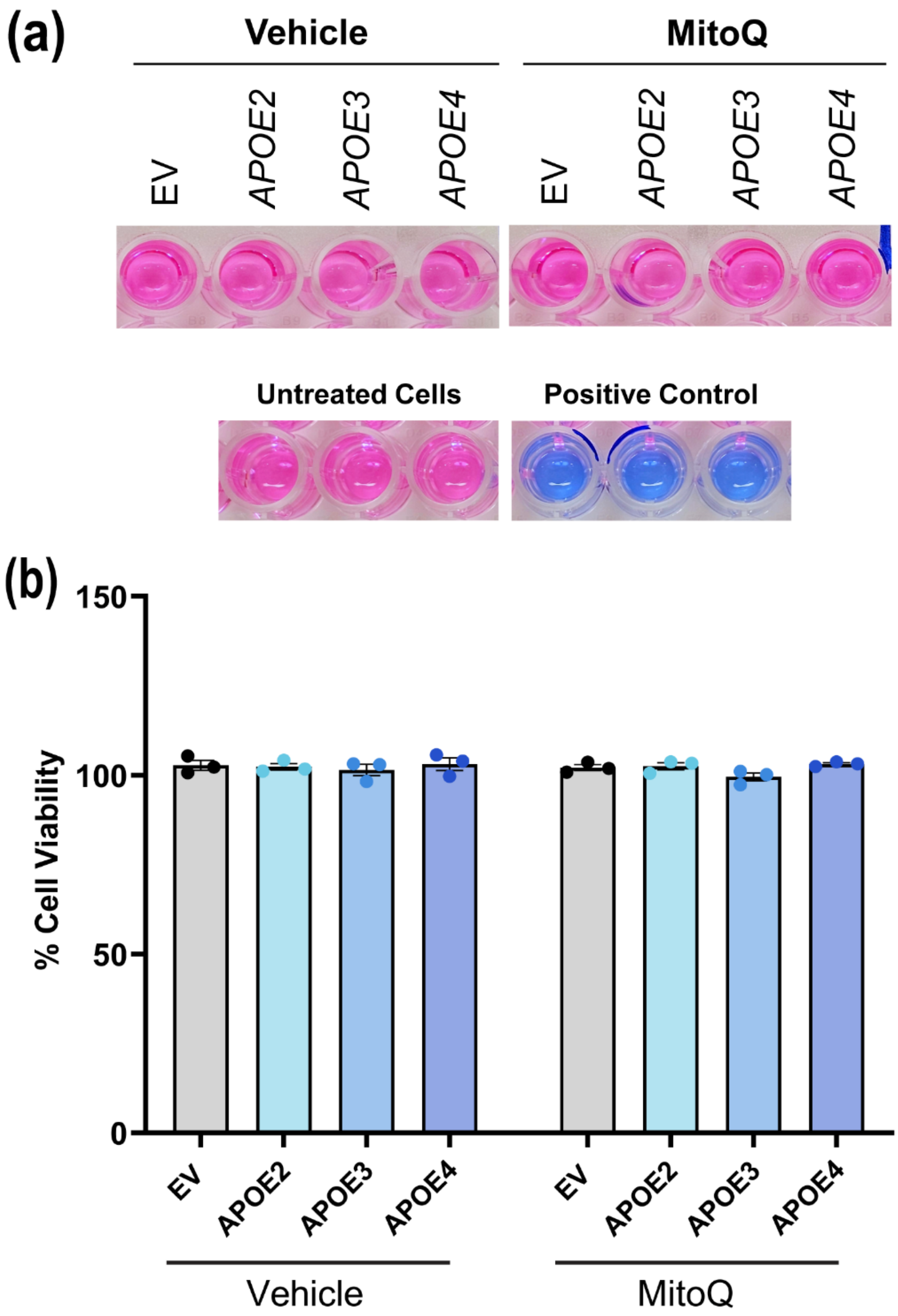


**Supplementary Figure 3: MitoQ does not affect cell viability in rMC-1.** (a) Representative images of Alamar Blue treated rMC-1 transfected with EV/ *APOE2*/ *APOE3*/ *APOE4*. Untreated cells and 20% DMSO-treated cells were used as control. (b) Bar graph showing quantification of % of cell viability, showing that 1μM MitoQ treated rMC-1 are viable compared to vehicle. Values are expressed as mean ± SEM (n: 3 independent experiments).

### Western blots for Figure 3

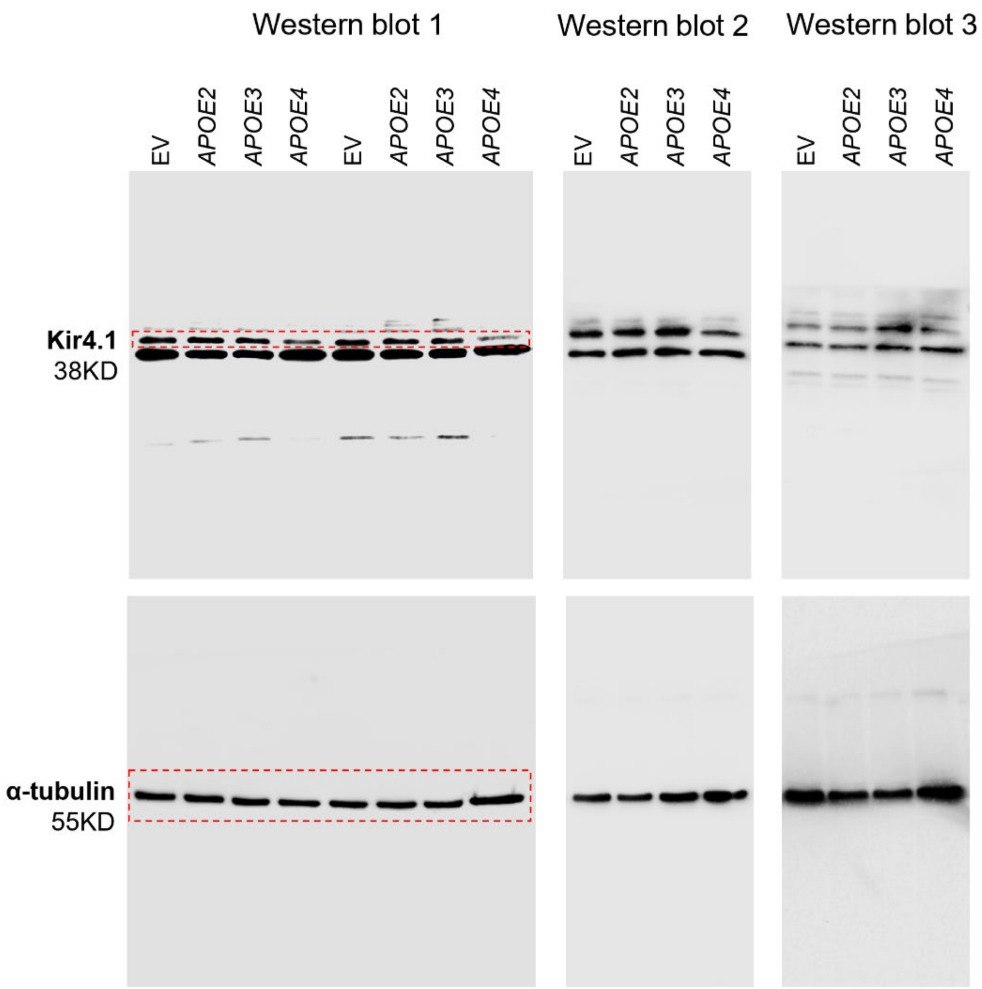

**Western blots for Figure 7**

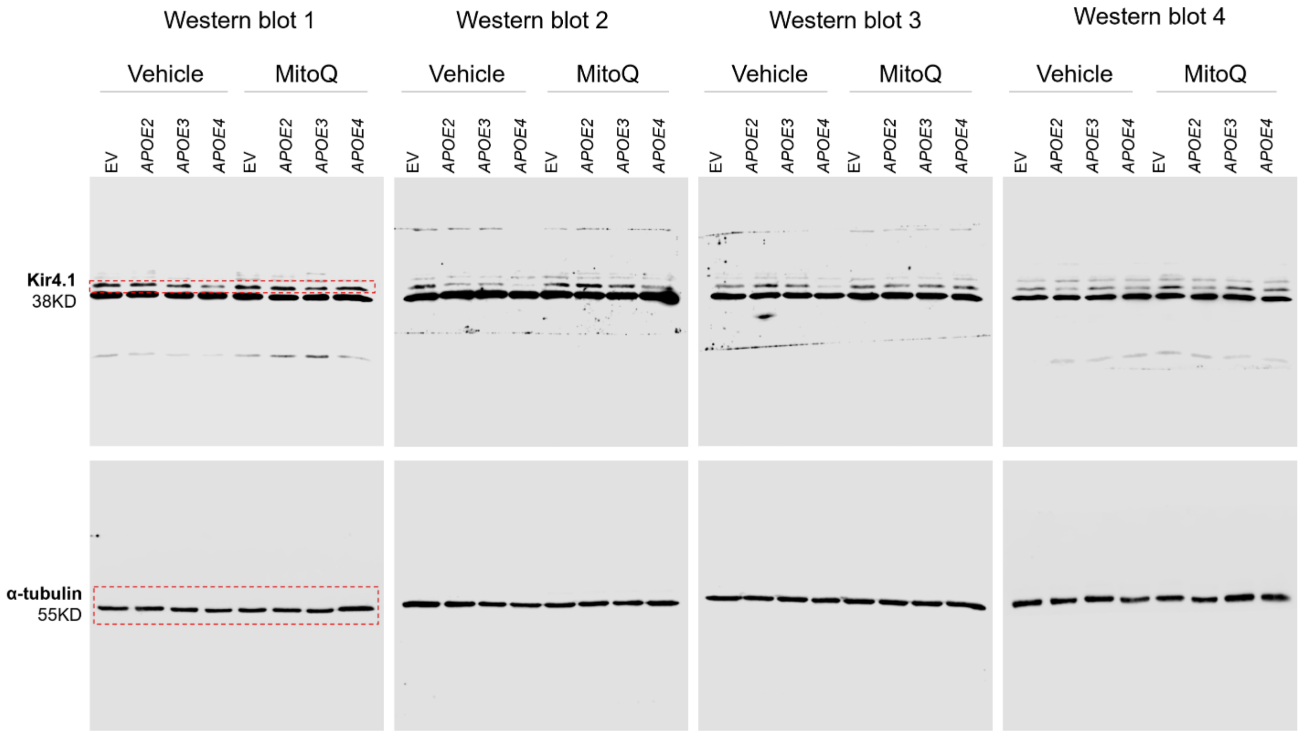
